# Acoustic Monitoring of Tropical Bats: *Saccopteryx bilineata’s* response to habitat and time across a restoration gradient using a 2D-CNN

**DOI:** 10.64898/2026.09.28.755061

**Authors:** Isabelle Secord, Xiuhan Zhang, Rhys Preston-Allen, Cristina Banks-Leite

## Abstract

Global biodiversity loss is a consequence of habitat degradation and fragmentation caused by the expansion of the Anthropocene and the overexploitation of natural resources. Passive acoustic monitoring enables longitudinal assessment of biodiversity but generates massive datasets. Advancements in machine learning have enabled the development of convolutional neural networks capable of automating this process. We developed a lightweight 2D convolutional neural network capable of detecting echolocation calls of the greater sac-winged bat, *Saccopteryx bilineata*, an insectivorous aerial edge-foraging bat, across a tropical restoration gradient in Pará, Brazil. The 2D-CNN model achieved a 97.84% accuracy and 100% precision. Deploying this model across 157,335 field recordings across 29 sampling points resampled over 3 years (2023-2025) yielded 2,098 positive detections of *S. bilineata*. Generalised linear mixed models revealed that time since reforestation significantly increased the odds of detecting *S. bilineata* (β = 0.22, p < 0.001), and that detection probability was 2.14 times higher in restoration plots than in forest habitats (β = 0.76, p = 0.0701). Acoustic hardware had a significant covariate influence in both fitted models (β = 0.52, β = 0.72, both p < 0.001). These findings demonstrate the practical applications of deep learning and bioacoustics to quantify habitat recovery and the value of restoration in regenerating functional ecological niches for aerial insectivores during early successional stages.

## Introduction

Habitat loss from land-use change and deforestation is the primary driver of biodiversity loss worldwide (Wiebe & Wilcove, 2025). A biome under particular threat is the Amazon rainforest in Brazil, home to over 10% of Earth’s terrestrial life, yet it has already lost nearly 13% of its original forest area (Flores et al., 2024). Deforestation disrupts ecosystem services, including water provision, carbon cycling, and climate regulation (da Cruz et al., 2021; Favretto & Hirota, 2026). The loss of these forests threatens wildlife populations with extinction and diminishes key ecological functions such as pollination, seed dispersal, and pest control (Wiebe & Wilcove, 2025). While active reforestation provides a promising solution, the timeframe required for wildlife to recolonise restored sites and recover ecological functionality remains poorly understood, largely due to insufficient long-term biodiversity monitoring.

Restoration in the tropics is an established nature-based solution that has gained significant traction through international initiatives such as the Bonn Challenge, which aims to restore 350 million hectares of degraded and deforested landscapes by 2030 (Dockendorff et al., 2022). Replanting native tree seedlings can directly counter historic forest loss across the Amazon’s ‘Arc of Deforestation’ in eastern Brazil (Nunes et al., 2020). Monitoring faunal recovery can be labour-intensive and expensive, and traditional survey methods such as point counts and mist-nets can underestimate or miss cryptic species. Focusing on a bioindicator target taxon, such as bats, offers a robust assessment of restoration success due to their diverse trophic guilds and sensitivity to habitat changes (Aodha et al., 2018; Cunto & Bernard, 2012).

Bats represent the second-largest order of mammals, comprising over 1,500 species globally with peak diversity in the tropics (Yovel, 2025). They are ideal targets for passive acoustic monitoring (PAM) because insectivorous species vocalise continually while echolocating, producing species-specific ultrasonic pulses suitable for automatic classification (Zualkernan et al., 2020). PAM provides a non-invasive, cost-effective method for tracking bat activity across space and time using autonomous recording devices (Silva & Herrera, 2026). However, the manual validation of bioacoustics data is time-consuming; convolutional neural networks (CNNs) offer an automated, scalable solution (Schwab et al., 2023; Silva & Herrera, 2026; Zualkernan et al., 2020). By converting raw audio into time-frequency representations (spectrograms) via the Short-Time Fourier Transform (STFT), 2D-CNNS can extract visual acoustic features to accurately classify echolocation calls (Alipek et al., 2023).

Despite these technological advances, little is understood about how tropical insectivorous bats utilise regenerating habitats during early-stage reforestation. Using deep learning within a PAM pipeline, this study evaluates the spatio-temporal response of *Saccopteryx bilineata* across a restoration gradient in the eastern Amazon. Specifically, this study addresses two research aims and a methodological objective:

• Aim 1 – Temporal response: Quantify the effect of time since reforestation on the binary detection probability of *S. bilineata* across restoration sites.

• Aim 2 – Spatial response: Compare the occupancy and detection probability of *S. bilineata* across habitat types by contrasting regenerating restoration plots with mature baseline forest stands.

• Methodological objective: Develop, train, and validate a lightweight 2D convolutional neural network optimised for accurate species identification within a large PAM dataset.

## Method

### Study System

Passive acoustic monitoring was conducted at an active restoration site covering 2890.5 hectares in the north-eastern state of Pará, Brazil (IBRD World Bank, 2024)(Fig. 1). Pará is the second-largest state in Brazil by area (∼125 mHa) and has over 94 mHa of natural forest, covering 76% of its land area (Global Nature Watch, 2026; Potapov et al., 2022). The average daytime temperature in Pará is 27 °C, with a nighttime average of 23 °C, resulting in a hot and tropical climate year-round (Climate: Pará in Brazil, 2026). Greater rainfall is experienced in the 1^st^ half of the year (Jan to May, average of 8.6-12.7 mm per day) than in the 2nd half of the year (July to Dec, average of 2.8 –5.9mm per day), with March being the wettest month and September the driest (Climate: Pará in Brazil, 2026).

**Figure 1:**
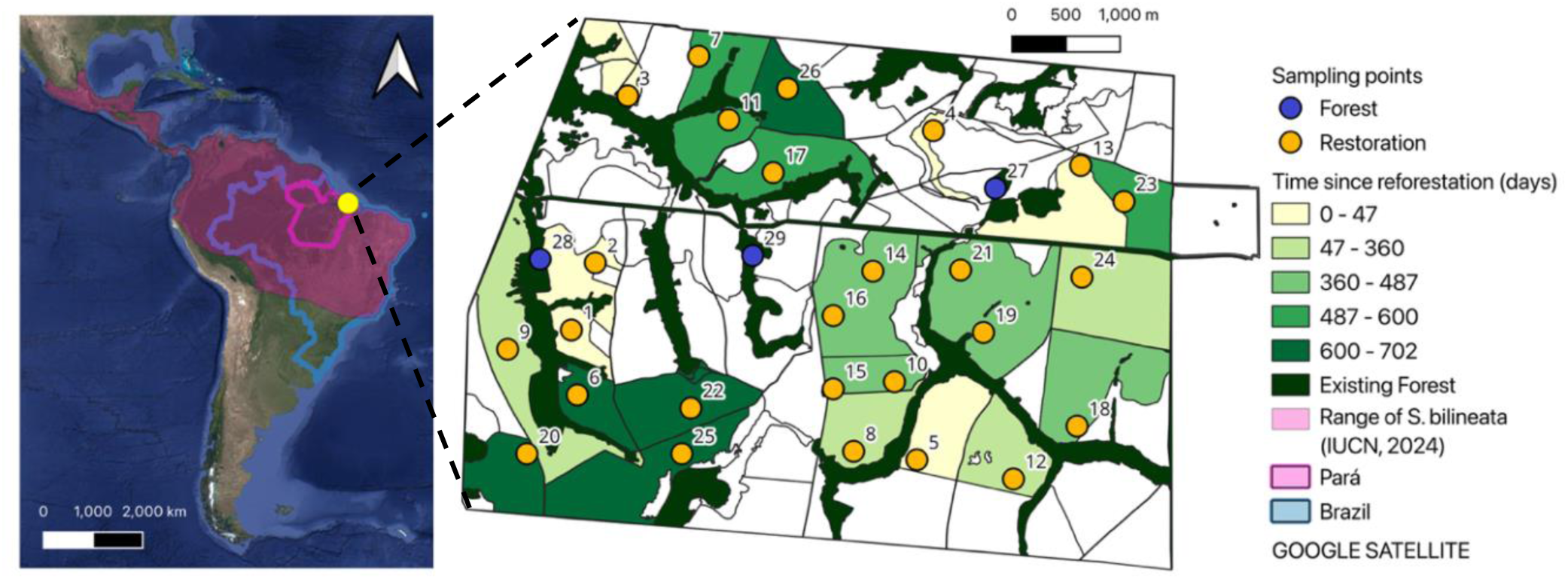
Map of South America showing the spatial range distribution of Saccopteryx bilineata in pink (IUCN, 2024). Brazil is outlined in blue, and the state of Pará is outlined in purple (IGBE, 2026; MapBiomas Brasil, 2026). A zoomed-in view of the study site is outlined by a black perimeter, with coloured areas in shades of green indicating the age of reforested areas and the locations of sample points. Light yellow areas were replanted more recently than the green areas, with dark green showing existing forest stands. Orange sample points were in restoration plots, and Blue sample points were in existing forest stands. Image produced in QGIS (v3.44 Solothurn).

Sampling was conducted over 7-10 days across three years, starting in February 2023, March/April 2024, and March 2025 (STable 2). There are 29 sampling points in total, spaced at a minimum of 500m apart – 26 were distributed across pastureland prior to restoration, and the remaining 3 were in existing forest stands (Fig. 1). The site is owned by Mombak, a carbon removal company, that is replanting trees using a seed mix of native and fast-growing species to maximise carbon sequestration, and has planted over 3 million trees in the state of Pará since April 2023 (Agence France-Presse (AFP), 2025; Ben Payton, 2025; Eduardo Laviano, 2023; MOMBAK, 2026). After 3 years of restoration, the tree saplings planted in the oldest restoration plots have grown to several metres in height (Agence France-Presse (AFP), 2025).

### Study Species

The greater sac-winged bat, *Saccopteryx bilineata*, is part of the Emballonuridae family of bats, and its geographical range extends from northern Mexico to across central Brazil (López-Baucells, 2018)(Fig. 1). Individuals weigh between 7 - 9 grams, with females typically weighing up to 15% more than males (Voigt et al., 2008). This bat is polygynous, with male bats defending territories for harems of four to eight females. *Saccopteryx bilineata* roost on external tree trunks and buttresses, with a single tree often supporting several harems and even other bat species, with each male *S.bilineata* defending an area of a few square metres (Altringham, 2011; Yovel, 2025). *S. bilineata* has dark dorsal fur, with two characteristic white stripes on its back and sac-shaped glands near its shoulders (Fig. 2b). *S. bilineata* has been well studied for exhibiting vocal babbling behaviour in pups (Fernandez & Knörnschild, 2017; Knörnschild et al., 2006; Knörnschild, Nagy, et al., 2012) and their complex courtship displays (Behr & von Helversen, 2004), which encompass acoustic, visual and olfactory cues from secretions, pheromones, and urine stored in their wing sacs (Fig. 2a)(Behr et al., 2009; Knörnschild, Nagy, et al., 2012; Voigt et al., 2008; Yovel, 2025). *S. bilineata* was assessed by the IUCN Red List as of least concern, but its population trends remain uncertain (IUCN, 2025).

**Figure 2a.**
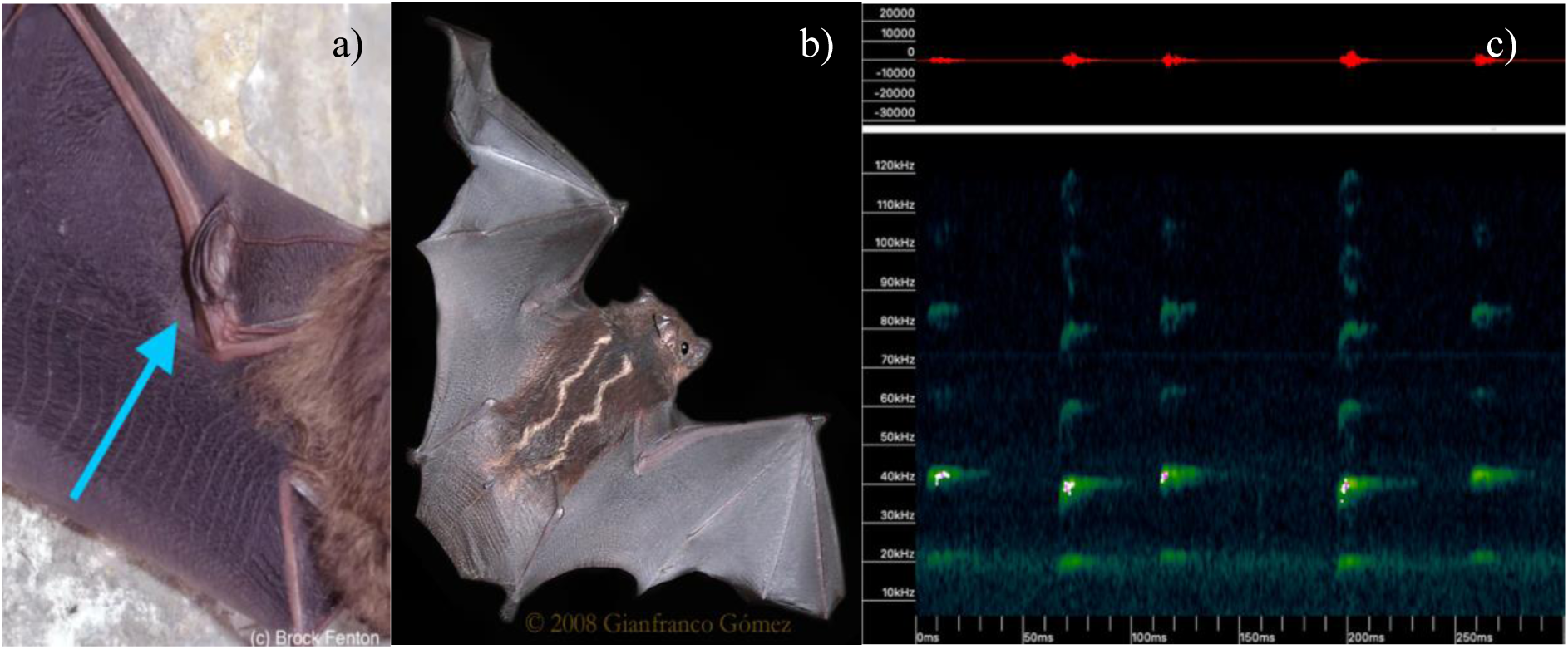
Blue arrow pointing at the sac pouch on the wing © Brock Fenton (Carter, 2012). 2b) A photo of Saccopteryx bilineata pelage and colour, with two dorsal white stripes along its back, © Gianfranco Gómez. (Gianfranco Gómez & Tracie Stice, 2009). 2c) Echolocation call showing alternating 43khz followed by 45khz, with harmonics visible. The 2^nd^ harmonic has the greatest energy. (Kaleidoscope 5.8.0.a)

*S. bilineata* is an aerial hawker and forages for insects along vegetation edges and open forest fragments, and can adapt its echolocation calls depending on vegetation clutter (Ratcliffe et al., 2011; Salazar-Pérez & Estrada-Villegas, 2025). This bat utilises an alternating echolocation call pattern comprising a lower-frequency pulse at ∼42kHz, followed by a higher-frequency pulse at ∼45kHz (Fig. 2c)(Burchardt et al., 2019; Jung et al., 2007; López-Baucells, 2018; Ratcliffe et al., 2011). Echolocation calls of this bat have been widely studied (Arévalo-Cortés et al., 2024; Arias-Aguilar et al., 2018; Jung et al., 2007; Knörnschild, Jung, et al., 2012; Ratcliffe et al., 2011), and the inter-pulse duration and the characteristic “chip chop” duplet pattern create a unique spatial signature that can be used by machine learning for a species-specific acoustic detection model (Alipek et al., 2023).

### Data Collection

Passive acoustic monitoring was conducted using two bat detector models due to equipment availability constraints: AudioMoths (v1 and v2, Open Acoustic Devices, 2017) and Song Meter Mini Bat units (v1.0.0 Wildlife Acoustics Inc., 2024). AudioMoths were deployed throughout 2023 and 2024, whereas Song Meters were co-deployed at a subset of sites in 2024 and used exclusively in 2025. Devices were mounted 1-2 m above the ground and orientated horizontally or slightly upwards to optimise capture of distant calls, with microphones pointed away from slopes and tree trunks to reduce acoustic interference (Fig. 3).

**Figure 3.**
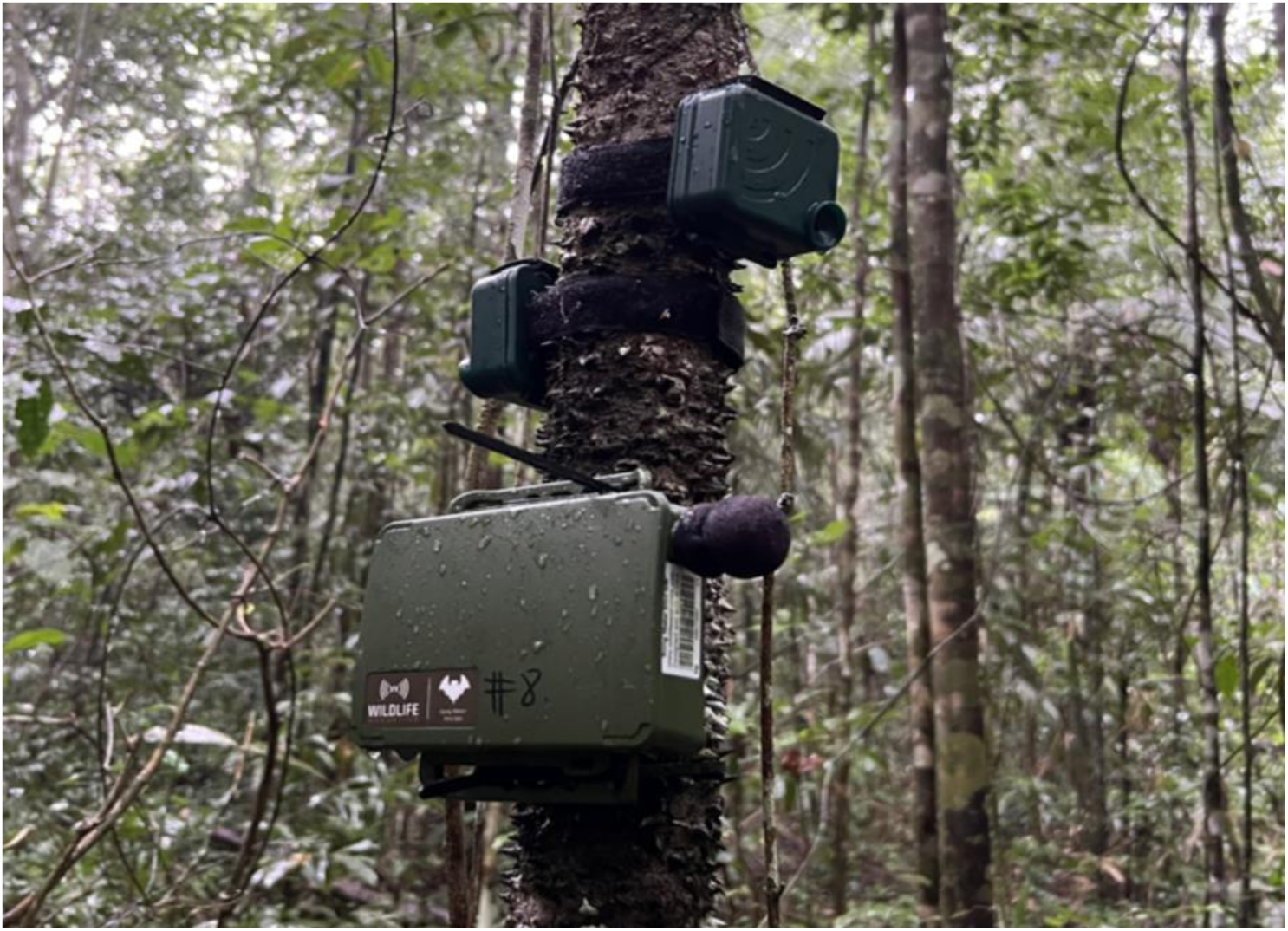
Devices co-deployed at a sample point. AudioMoth device positioned at the bottom, with a black microphone visible. With two Song Meter devices deployed above, the left device set to capture audible calls and the right device set to capture ultrasonic frequencies © Photo taken by Rhys Preston-Allen

All devices recorded at a sampling rate of 256 kHz, capturing frequencies up to 128 kHz. Duty cycling and recording schedules differed between device models. AudioMoths were scheduled for continuous duty cycling (1-minute recording every 5 minutes from 1 hour before sunset to 1 hour after sunrise (Kloepper et al., 2016)). Song Meters were set to ultrasonic triggering, recording audio clips up to a maximum duration of 15 s when triggered by signals exceeding the minimum threshold frequency of 16 kHz.

### 2D-CNN detection model summary

To detect *Saccopetryx bilineata* calls in the wider passive acoustic monitoring dataset, the architecture of an existing 1-dimensional convolutional neural network (1D-CNN) (*BruceWayne,* Canosa, 2025) was adapted to a 2-dimensional convolutional neural network (2D-CNN). The 2D-CNN model was built using *Keras* (v3.14.1, Chollet, 2015) and *TensorFlow* (v3.14.1, TensorFlow Developers, 2026) in *Python* (v3.11.9). Deep learning models like CNNs can recognise and extract meaningful patterns from images, such as echolocation pulses in spectrograms, to detect and identify bat sounds from background noise (Alipek et al., 2023; Aodha et al., 2018, 2022). This model uses a 2-dimensional approach to scan the spectrogram across time (x-axis) and frequency (y-axis) to learn the spatial signature of *S. bilineata’s* alternating echolocation.

### Raw Audio Preprocessing

All audio files were pre-processed with *librosa* (v0.11.0) to generate consistent input feature tensors for model training and detection using a preprocessing function defined by Canosa (2025). Recordings were sampled at 256 kHz and divided into non-overlapping segments of 0.5s (128,000 samples), with longer clips truncated and shorter clips zero-padded to maintain the correct segment length. A window size of 0.5s with no overlap was chosen to capture at least two sets of alternating pulses, as the average inter-pulse window was 100-200ms, with a within-pulse duration of 50ms between the alternating frequencies (Jung et al., 2007; Knörnschild, Nagy, et al., 2012; López-Baucells, 2018) (Fig. 2c, Fig 4). Segments were Short-Time Fourier transformed (STFT) and mapped onto the Mel scale across 256 bins, at a frequency range of 20 kHz to 100 kHz to capture the target ultrasonic range. The power spectrogram was converted to decibels and dynamically scaled by the maximum value. Lastly, the inputs were transposed to produce a 2D tensor with shape 251 time steps x 256 Mel bins, ready for feature extraction by the 2D-CNN.

**Figure 4.**
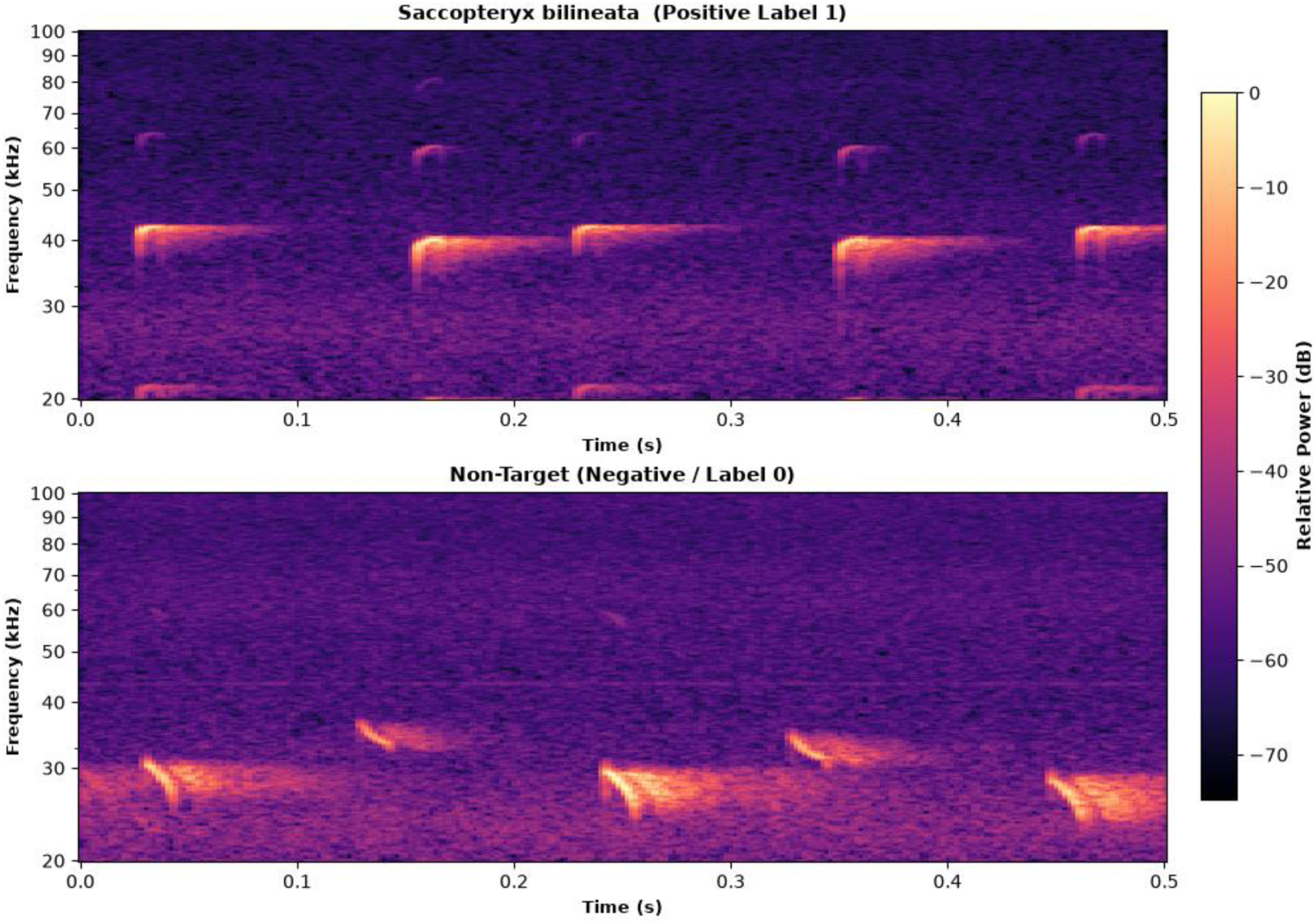
A comparison of two spectrograms of a positive (1) and a negative (0) recording from the training library. The top panel shows an example of an Saccopteryx bilineata echolocation call with the species-specific alternating-frequency pulses that form the spatial signature the automated classifier learns. The bottom shows a non-target bat call. Code provided by Canosa, 2026.

### Training Library

A training library of 916 clips was compiled from an annotated subsample of the dataset with annotations provided by a bat acoustic expert in the lab group. Parent audio files were preliminarily inspected visually and audibly in Kaleidoscope Lite (v5.9.0a) to confirm bat presence. To build a standardised library, an interactive Python script used *soundfile* (v0.14.0) and *librosa* to resample audio to 256 kHz and locate the root-mean-square (RMS) energy peak in the file. Candidate windows were visually inspected using a 256-mel-bin spectrogram to confirm the capture of clear *Saccopteryx bilineata* echolocation pulses, after which they were saved as a 0.5-s clip (Fig. 4). Files containing non-target bat calls and/or ambient background noise were saved as negative clips. Only one clip was collected from each parent file to prevent data leakage in training.

The resulting library contained 707 negative clips and 209 positive clips, yielding a training-class imbalance ratio of 3.55:1. To preserve this ratio, the library was randomly stratified into 3 subsets: training (70%), validation (15%), and test (15%). Model performance was monitored over learning epochs using standard performance metrics, including accuracy, precision, recall, binary cross-entropy loss, area under the receiver operating characteristic curve (AUC-ROC) and a confusion matrix (Rainio et al., 2024; Sharma et al., 2023)(S Table 1).

### Model Training And Architecture

To account for the training-class imbalance, a custom binary loss function (Lin et al., 2018) was implemented and adapted from Canosa (2025). The weighting factor (Alpha, α=0.78) scales greater loss penalties to the positive class and dampens weights from the negative classes, whilst the focusing parameter (gamma, γ=2.0) directs the model to concentrate on harder examples and reduce weights for easy examples. The *AdamW* optimiser (learning rate = 0.0001, weight decay = 0.001) was used for decoupled weight decay (Loshchilov & Hutter, 2019), and an *L2 regularisation* penalty (λ=0.0001) was implemented to prevent overfitting. *ReduceLROnPlateau* (Mukherjee et al., 2019) reduced the learning rate by a factor of 0.5 if it did not improve for 20 epochs, with a minimum rate of 0.000001.

Early stopping prevented the model from overfitting and initiated monitoring from epoch 20, to restore the best weights if the minimum improvement (Δ = 0.001, patience threshold = 30 epochs) was not met. Model checkpoints were saved based on the maximum validation AUC score. Training was set for a maximum of 200 epochs, with accuracy, precision, recall, and AUC evaluated by *keras.metrics (Keras, 2015)*.

The 2D-CNN architecture consisted of four feature extraction blocks (Fig. 5) with feature-scale units of 32 -> 64 -> 128 -> 128. Each block contained a 2D convolutional layer with 3×3 kernels, a *Rectified Linear Unit (ReLU)* activation function, and L2 regularisation. Batch Normalisation and 2×2 Max Pooling followed the 2D convolution layer. Extracted spatial features were then flattened before passing through a dense layer with 128 units, with ReLU activation function and L2 regularisation, and a dropout layer of 0.4 before terminating in a sigmoidal activation function to return a binary classification. Following training, the saved model’s optimised threshold was set to t=0.95.

**Figure 5.**
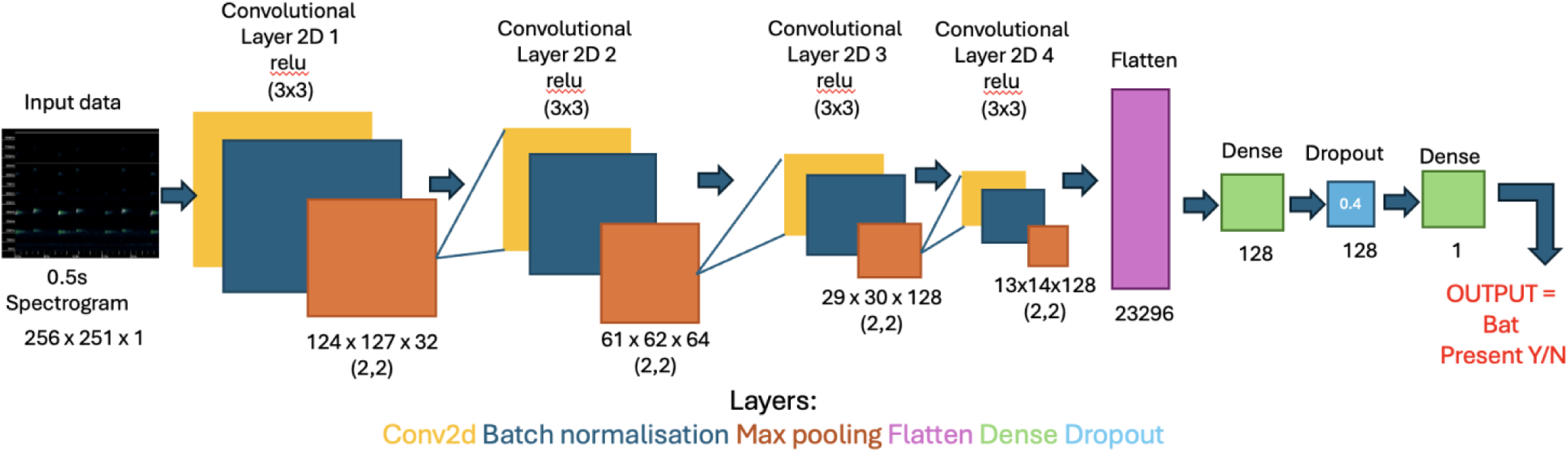
Visual representation of 2D-CNN model, input is a 0.5s spectrogram, output is a confidence score resulting in a binary classification of Yes or No to indicate if the target echolocation calls were present in the clip. The convolutional layers decrease over 4 subsequent layers before being flattened and passed through a dense layer with dropout of 0.4, followed by a final dense layer. The threshold for positive classification is 0.95. See the appendix for the flowchart of tensor shapes SFig 13, SFig 14.

### Model Validation

The wider acoustic data set was analysed for *Saccopteryx bilineata* calls using the 2D-CNN. A stratified validation approach was used to assess baseline model performance and the performance at the threshold boundary (t=0.95) across all sites and years. A total of 709 files were audited and grouped into five confidence tiers. Audit tier 1 grabbed high-confidence (>0.98) files to confirm true positives, tier 2 grabbed borderline positives (0.95<x<0.98), and tier 3 grabbed borderline negatives (0.90<x<0.95) to validate files on either side of the 0.95 threshold. Tier 4 grabbed lower confidence files (0.80<x<0.90) to confirm no faint target calls were incorrectly classified as negatives, and tier 5 grabbed very low-confidence files (<0.20) to confirm true negatives. All files were manually validated in their entirety by visual and audible inspection in Kaleidoscope Lite (v5.9.0a).

### Explanatory Variables And Statistical Analysis

The primary response variable was the binary detection of *Saccopteryx bilineata* within individual acoustic recordings (1 = present, 0 = absent). A recording file was classified as present (1) if it contained at least one high-confidence (>0.95) 0.5s segment identified by the automated detection model. Device recorder type (Song Meter or AudioMoth) was included as a fixed control covariate across all models to account for systematic differences in recording sensitivity and deployment schedules. The proportion of positive recordings for a sample point was calculated as the number of recordings in which *S. bilineata* was detected, divided by the total number of recordings, and used as a measure of naïve detection probability to enable comparison across sample points and years.

Habitat was treated as a binary explanatory variable, based on the immediate environment surrounding each sample point. Sample points within existing forest stands that predate the reforestation efforts were classified as “Forest”, while pasture and active restoration sites were both classified as “Restoration”. Time since reforestation was quantified as a continuous explanatory variable, measured in days. This was calculated as the difference between the date of the specific acoustic survey night and the documented replanting date for each sample point (time since reforestation (days) = Survey night date - Replanting date).

To investigate the factors influencing the binary detection probability of *Saccopteryx bilineata* in the wider acoustic dataset, separate generalised linear mixed models (GLMM) were fitted for each research aim. To facilitate model convergence and standardise effect sizes, time since reforestation was z-standardised (mean = 0, SD = 1) prior to modelling. For aim 1, forest age (z-scaled) and device type were included as fixed effects; for aim 2, habitat type and device type were included as fixed effects. In both models, the sample point was included as a random intercept to account for repeated measures across the sampling period (2023–2025). Models were fitted with a binomial error distribution and a logit link function (family = binomial).

All statistical analyses, data manipulation, calculation of explanatory variables, and plotting were conducted in R (v 4.4.1) using RStudio with the following packages: *DHARMa (v0.5.0 Hartig, 2026), dplyr (v1.2.0 Wickham et al., 2026), ggeffects (v2.3.2* (Lüdecke, 2018)*, ggplot2 (v4.0.2 Wickham, 2016), lme4(2.0.6 Bates et al., 2015), performance(v0.17.1* Lüdecke et al., 2021)*, tidyverse (2.0.0 Wickham et al., 2019)* and *viridislite (0.4.3 Garnier et al., 2024)* .

## Results

The 2D-CNN model analysed the wider acoustic dataset (n=157,335 recording files, ∼1,008 hours) for *Saccopteryx bilineata* echolocation calls across 29 sampling points over three monitoring years (2023-2025)(Fig. 6). *Saccopteryx bilineata* was detected in 2,098 recordings, corresponding to an overall dataset detection rate of 1.33% at a model confidence threshold of 0.95. Across the survey, the number of positive detections per point ranged from 0 to 177 (median = 11, IQR =19).

**Figure 6.**
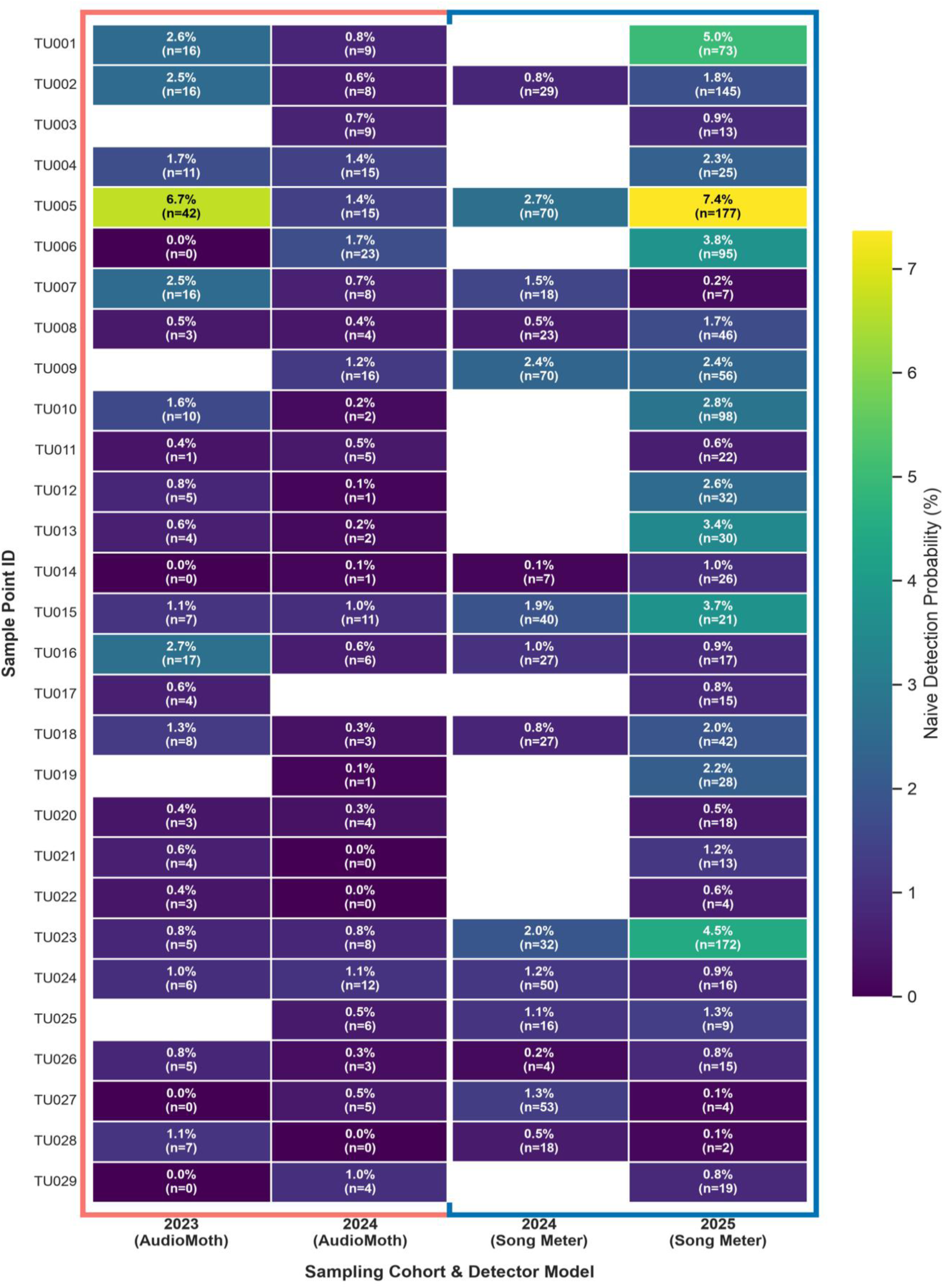
A heatmap summary table showing the sampling point IDs along the Y axis and the sampling cohort years and recorder type across the X axis. Within the cells are the naïve detection probability and the number of positive calls, n. The heatmap is colour-coded to show points with a lower positive detection rate in deep purple and sites with a higher positive detection rate in blue, progressing to yellow-green. The pink outline indicates the first two sampling years used AudioMoths, and the blue outline indicates that a few sample points in 2024 and all in 2025 used Song Meters

At the sample-point level, the naïve detection probability ranged from 0.00% to 7.37% across the study period, with an overall deployment median of 0.81% (IQR = 1.21%). In 2023, the sample-point detection median was 0.79% (IQR of 1.20%). In 2024, the two devices yielded different detection profiles: AudioMoths reported a median of 0.49% (IQR 0.62%), whereas the Song Meters reported a median detection probability of 1.10% (IQR 1.06%). In 2025, when Song Meters were deployed exclusively, the median sample-point probability increased to 1.34% (IQR of 1.74%).

### Model Assessment And Validation

The 2D-CNN demonstrated high discriminative capability (F1 = 0.9647) when evaluated on an unseen, held-out test dataset (n=139; non-target clips = 95; positive target clips = 44). At the decision threshold of 0.95, the model achieved an ROC-AUC of 0.9983, an overall test accuracy of 97.84%, 100% precision (zero false positives) and a recall of 93.18% (3 false negatives) (SFig. 9) During training, the model converged to an optimal checkpoint at epoch 22 (SFig. 10), achieving a validation loss of 5.41%, a validation precision of 100%, a validation accuracy of 98.52%, and a ROC-AUC of 1.000.

Stratified manual validation of 709 files assessed model performance across five confidence tiers spanning either side of the 0.95 decision threshold. Tiers 1 (>0.98) and 2 (0.95 < x < 0.98) classified recordings as positive, and manual spectrogram verification of 314 files in Kaleidoscope Lite yielded a true-positive precision of 43.63% across both tiers (true positives = 137, false positives = 177). Tier 1 achieved a precision of 52.17%, while Tier 2 had a lower precision of 36.93%, due to false triggers from heavy rainfall and insect noise.

Tiers 3, 4, and 5 evaluated 395 files classified as negative, of which 324 were confirmed true negatives (82.03%). Tiers 3 and 4 evaluated files within the confidence range 0.80 <x < 0.95, of which 71 (17.97%) were false negatives containing faint *S. bilineata* echolocation calls partially masked by overlapping non-target bat calls. Manual verification of 82 low-confidence (<0.20) files in tier 5 yielded a true negative rate of 97.56%, with only two very faint, distant *S. bilineata* calls missed by the model. Across the full stratified audit (n=709), which specifically targeted borderline and baseline cases, the model achieved an overall detection accuracy of 65.02%, with a recall of 65.87% and an F1 score of 52.49%.

### Response To Time Since Reforestation

Data from 84 survey deployment windows (STable 2) across 24 sampling points with a documented replanting date contributed 133,318 recordings (∼844 hours), of which 1,899 contained positive *Saccopteryx bilineata* echolocation recordings. Time since reforestation had a significant effect on *S. bilineata* binary detection probability (β = 0.22, SE =± 0.049, z = 4.53, p < 0.001)(Fig. 7). For every additional year that a restoration plot of replanted trees matures, the odds of detecting *S. bilineata* increased by 43.5% (OR = 1.44, 95% CI = [1.23, 1.68]).

**Figure 7.**
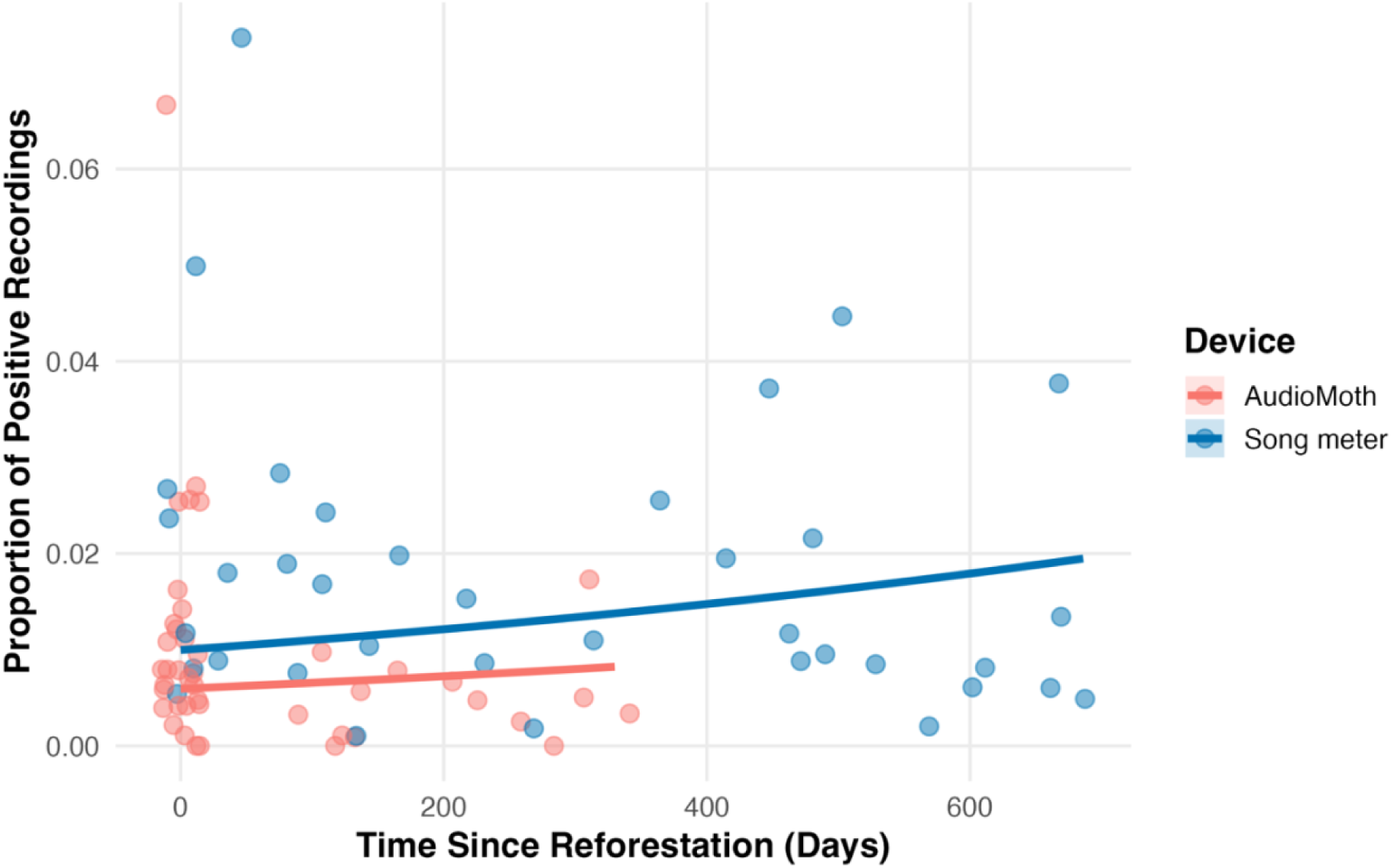
Proportion of positive Saccopteryx bilineata recordings as a function of time since reforestation (in days) Each point (n=84) represents an individual deployment across 24 sampling locations resampled across 2023-2025. AudioMoth recordings in pink, Song Meter in blue. Fitted lines show device-specific linear trends, with points jittered and with reduced opacity to illustrate sample density.

The model also showed that the hardware covariate had a significant influence on *S. bilineata* detection probability (β=0.52, SE±0.072, z=7.29, p < 0.001). The GLMM explained 19.3% of the total variance (conditional R^2^ = 0.193), and the random intercepts for sample points accounted for the spatial nesting and repeated-sample design (σ^2^= 0.65). Model convergence was facilitated by z-scaling forest age, and the DHARMa test confirmed an even residual distribution, with no overdispersion (p=0.912).

### Response To Habitat

The influence of habitat on the binary detection probability of S. bilineata was assessed across the entire acoustic dataset from 29 sample points over three years, comprising 157,335 recordings (∼1008 hours). A GLMM was fitted with hardware and habitat type as fixed effects and sample point as a random intercept. Detection odds of *S. bilineata* were 2.14 times higher in restoration points compared to existing forest stands (OR = 2.14, 95% CI = [0.94, 4.88]), representing a positive trend that approached but did not reach conventional significance (β=0.76 SE=±0.42, z=1.81, p=0.0701)(Fig. 8). Hardware also exhibited a strong and highly significant control-covariate effect (β=0.70, SE=0.06, z=11.80, p<0.001). The model accounted for 15.4% of the total variance (Conditional R^2^=0.154), and random intercepts for the sample point capturing site-level variance( σ^2^ = 0.44). DHARMa diagnostic tests confirmed adequate model fit with no overdispersion detected (p=0.888).

**Figure 8.**
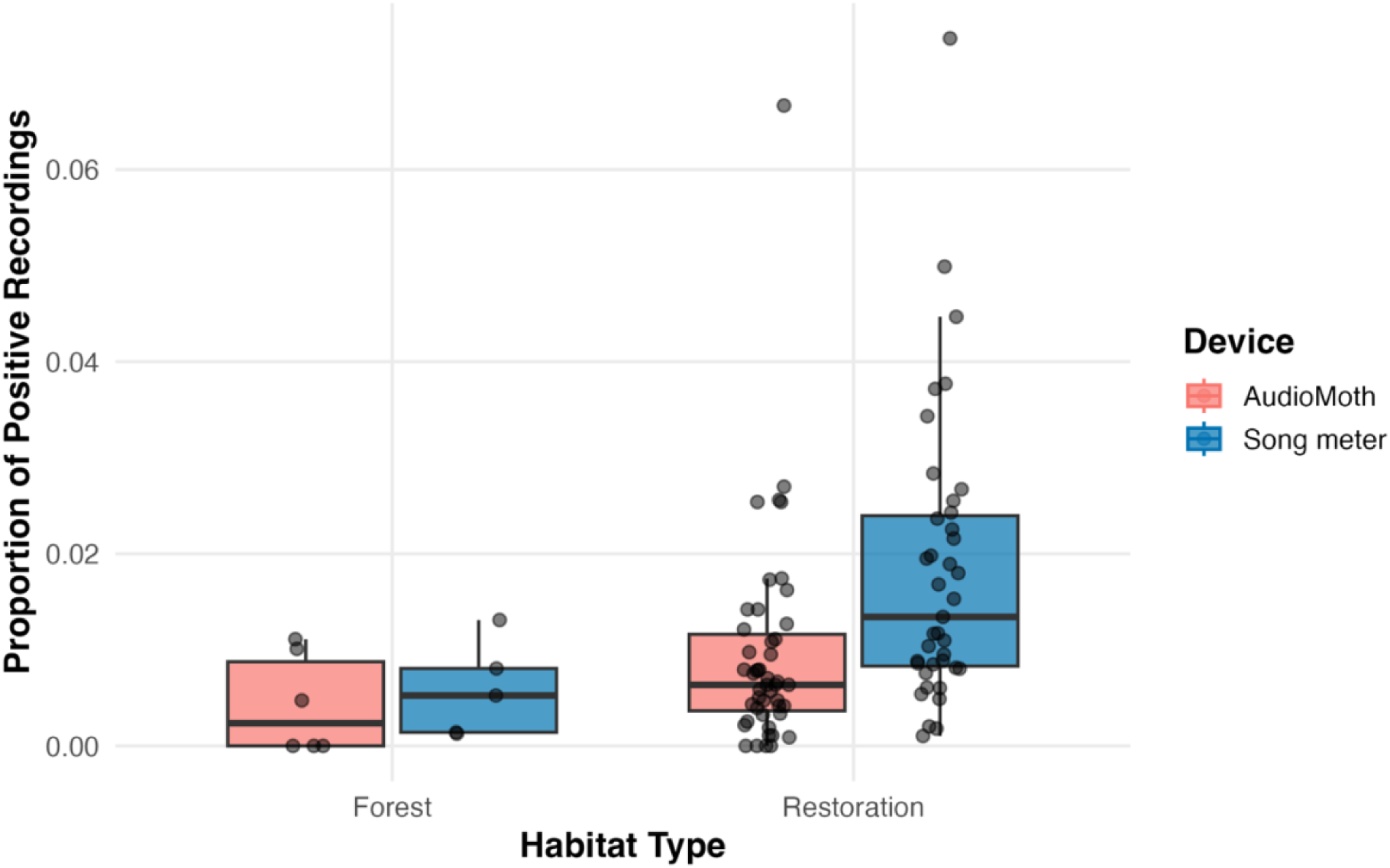
Proportion of positive Saccopteryx bilineata recordings across habitat types (Forest vs Restoration) with AudioMoth in pink and Song Meters in blue. Each point (n=101) represents an individual deployment across 29 sampling locations, resampled from 2023-2025 (11 forest deployments across 3 sites and 90 restoration deployments across 26 sites). The boxes show the interquartile range (IQR), the black horizontal line shows the median, and the whiskers extend to 1.5×IQR. Individual deployment points are jittered with reduced opacity to illustrate the sample distribution.

## Discussion

This study used a passive acoustic monitoring dataset and a deep-learning classification pipeline to investigate the spatiotemporal response of *Saccopteryx bilineata* to baseline forest stands and active reforestation plots across a restoration gradient. The acoustic detection and monitoring of bats provides an effective bioindicator framework (Aodha et al., 2018; Jones et al., 2009; Russo et al., 2018, 2021) for evaluating the success of restoration schemes and ecosystem recovery in post-agricultural landscapes. Three key findings were reported in this study.

Firstly, *Saccopteryx bilineata* exhibited a twofold (x2) higher detection probability in regenerating plots than in mature forest patches, contributing to a greater understanding of this bat’s foraging preferences and habits, as well as the functional value of successional edge habitats for aerial insectivorous bats. Secondly, as reforested plots matured over time, *S. bilineata* detection probability increased significantly (p<0.001, OR = 1.44, ), demonstrating that greater sac-winged bats successfully colonise early secondary forests and that more activity is recorded in established restoration locations over time. Finally, recording devices exerted a substantial and significant influence on the naïve detections across both GLMMs, highlighting the need to consider technological covariates in long-term bioacoustics monitoring datasets.

### Response To Habitat

*Saccopteryx bilineata* is an edge-space aerial insectivore (Denzinger et al., 2018), and our results found higher naive detection probabilities and higher odds of occurrence in active restoration plots than in existing forest stands. Although this relationship approached (β = 0.76, p=0.071) but did not reach statistical significance (p=0.05), the effect size was very large (OR = 2.14, CI = [0.94, 4.88]), indicating a strong ecological response consistent with the species’ biology. Bats occupy different foraging niches, and *Saccopteryx bilineata* hunts for flying insects in semi-open spaces above low-level shrub vegetation, capturing prey in flight (Denzinger et al., 2018; Jakobsen et al., 2015; Jung et al., 2007). During the pursuit of prey, *S. bilineata* produces a high-energy feeding buzz, generating a broad acoustic field of view with a wide sonar beam, but this strategy ensonifies a large spatial volume, creating echoes from nearby vegetation clutter that can mask prey (Jakobsen et al., 2012, 2015). Therefore, by hunting over open vegetation rather than in forest interiors, as seen in our results, *Saccopteryx* can avoid feedback echoes from clutter, maximising prey detection range and capture success.

Additionally, the bioacoustic adaptations of *S. bilineata* echolocations are specialised for navigating edge spaces. Jung et al. (2007) demonstrated that the species’ characteristic alternating frequency pulses (42kHz and 45kHz) enable them to discriminate between echoes of succeeding calls, making them adept at foraging along forest edges and preventing pulse-echo ambiguity. This sensory perception was further supported by Ratcliffe et al. (2011) who observed that *S. bilineata* dynamically adjusted call interval and sequence duration when approaching complex habitat edges to optimise spatial field of view. In addition, Salazar-Pérez and Estrada-Villegas, (2025) proposed that *S. bilineata* hunts in open and successional spaces to access prey more effectively via unobstructed flight paths and to reduce direct competition with clutter-tolerant forest-specialist insectivorous bats. In summary, the observed higher detections of *S. bilineata* in restoration plots reflect their specialised echolocation sequence and foraging niche, thereby validating these results.

### Response To Time Since Reforestation

Forest restoration can buffer the adverse effects of habitat loss and fragmentation on biodiversity (Rowley et al., 2024), and our models revealed a significant positive relationship between naïve detection of *S. bilineata* and time since reforestation (β = 0.22, p < 0.001). Age of restoration plots remained a robust predictor of activity even after controlling for the significant effects of recorder type (β =0.52, p<0.001), with the odds of detection increasing by 43.5% with every additional year of plot maturation (OR=1.44, CI = [1.23,1.6]). These findings align with Rowley et al. (2024) who observed that vegetation structure significantly predicted *S.bilineata* activity across a landscape matrix that contained a 30-year-old secondary forest. They also corroborate Denzinger et al. (2018) regarding vertical niche differentiation among sympatric emballonurids, where *S. bilineata* preferentially exploits low shrubs and edge strata, while *Saccopteryx leptura* forages higher in the subcanopy and canopy.

*Saccopteryx bilineata* is commonly observed roosting on tree trunks and buttresses and relies on vertical surfaces for harem territories (Altringham, 2011; López-Baucells, 2018; Yovel, 2025). As replanted saplings grow and mature over the study period (2023-2025), vertical surface area and tree girth will increase, thereby generating the structural architecture required for roosting and courtship displays (Behr & von Helversen, 2004; Voigt et al., 2008). Previous studies on *S. bilineata*’s response to vertical stratification and physical trapping with mist nets (Bernard, 2001; Yoh, Clarke, et al., 2022) confirmed that *S. bilineata* utilises both the higher and lower strata and exploits vertical open spaces for foraging.

Over time, the structural homogeneity of active restoration plots will change as agricultural pasture is replaced with young trees, transitioning into a heterogeneous mosaic that can support greater insect diversity and attract insectivorous bats (Marques et al., 2016). Jung et al. (2026) evaluated bat responses to rewilding interventions and found that time since treatment implementation resulted in higher feeding and echolocation activity among edge-foraging insectivores, directly supporting our results. Restoration efforts that create diverse canopies and vegetation structures will promote microclimatic variation and increase prey availability, thus restoring functional niches for tropical insectivorous bats (Wild et al., 2026).

### Model Performance On The Annotated Dataset

The 2D-CNN performed very well on the annotated dataset and demonstrated strong discriminative capability on the unseen annotated test dataset (n=139), achieving zero false positives and three false negatives at the 0.95 threshold (F1 = 0.9647, ROC-AUC= 0.9983). The model’s high F1 score aligns with findings by Sharma et al. (2023), who found that deep learning models trained on datasets with a broad representation of the complex environment captured greater variability in the target species, achieving higher F1 scores. Including ‘empty’ background noise clips and non-target bat calls across various frequencies in the training library fine-tuned the model’s ability to isolate the specific acoustic characteristics of *S. bilineata* calls from the ambient background soundscape and to ignore discrepancies (Silva & Herrera, 2026). Implementing standard regularisation and CNN architecture practices tailored for bioacoustics classification models (Stowell, 2022) optimised model performance and ability and prevented overfitting on the training cohort. (SFig. 9)

### Model Performance On The Non-Annotated Dataset

Evaluating the model’s true performance on 709 stratified non-annotated field recordings revealed an overall accuracy of 65.02% and a recall of 65.87%. As this audit specifically targeted and oversampled near-threshold files (0.80<x<0.95), the resulting precision (43.63%) reflects the challenges that automated classifiers encounter when processing heterogeneous real-world data. Environmental noise can interfere with and mask target echolocation signals, thereby increasing false detections (Zualkernan et al., 2020). Variation in echolocation calls, such as adapting search phase pulses into feeding buzzes during foraging (Walters et al., 2013), and the distance of the target vocalising species from the recording device (Aodha et al., 2018; Sharma et al., 2023) can also introduce variation in calls that are subsequently missed by the classifier. The quality and diversity of recordings, as well as the use of sufficiently large and varied training data, are paramount to deep learning networks succeeding on real-world datasets.

Nonetheless, despite these edge-case misclassifications, manual review of 157,335 field recordings would take over 126 eight-hour working days, given that the dataset contained 1,008.7 hours of raw audio. This data processing bottleneck is greatly reduced by computer automation, and deploying the 2D-CNN model on the wider dataset via parallel processing on a high-performance computing cluster took only a few hours (Harvey, 2017). Using a strict operational threshold (t = 0.95), combined with sample-point random intercepts, ensured that residual classification uncertainty did not undermine the broader ecological relationships. (SFig. 11)

### Comparison Of The Model

The architecture, training efficiency, and discriminative performance of the 2D-CNN developed in this study align with recent advances in bioacoustics and support the adoption of lightweight convolutional networks for targeted bat monitoring. Alipek et al. (2023) developed a 3-layer supervised CNN that achieved high predictive accuracy for German bat recordings captured at different heights and spatial locations, and found the simplicity of the network still provided high predictive capabilities on the dataset, illustrating that deep architectures are not necessary for bat classifiers. Whilst large datasets can improve CNN performance, Yoh et al. (2022) reported that expanding the training dataset for their Borneo bat classifier (BBC) from 1000 to 2000 or 5000 calls provided diminishing returns in accuracy (>85%) relative to the substantial increase in subsequent computational power. Our 2D-CNN model achieved a high F1 score and accuracy when trained on fewer than 1000 annotated clips (n=916), demonstrating the utility of a targeted architecture for lightweight models and strong performance on modest training samples.

The lightweight design of the 2D-CNN offers computational advantages over deep architectures and heavy multi-species classifiers. Kobayashi et al. (2021) developed a 30-species acoustic classifier for Japanese bats that achieved an identification accuracy of 98.1% but required 877.8 hours to train, likely due to the network depth being overpowered. Conversely, our dedicated *S. bilineata* model converged to an optimal checkpoint in just over an hour on standard computing hardware. With 916 training clips and its lightweight architecture, our 2D-CNN successfully learned features of the target call in just four layers.

Additionally, differences in acoustic hardware across multi-year studies present a challenge as data outputs, recording regimes, and microphone sensitivity can affect model generalisability. Detection probabilities of *Saccopteryx bilineata* were significantly different between the two devices, AudioMoths and Song meters (aim 1 GLMM:β = 0.52, p < 0.001; aim 2 GLMM:β = 0.70, p < 0.001), but the 2D-CNN still successfully identified target calls in field recordings from both devices. Schwab et al. (2023) demonstrated that CNNs that are sufficiently deep for the task can robustly handle disturbances from environmental or technical noise, even when field recordings use equipment that differs from the devices used to collect the calls in the training database. Finally, our findings corroborate Aodha et al. (2022) and their results from their BatDetect2 pipeline, which evaluated a dataset of 285 passive acoustic recordings from the Yucatan Peninsula. Aodha et al. (2022) achieved high precision-recall performance for *Saccopteryx bilineata*, using 381 training calls and 92 test calls, demonstrating that the distinctive alternating frequency echolocation pattern of *S. bilineata’s* calls provides a highly learnable spectrogram signature. The success observed in these studies demonstrates that convolutional neural networks can reliably identify *Saccopteryx bilineata’s* presence, even in acoustically complex tropical soundscapes.

### Limitations

One limitation of this study is the unbalanced survey effort across habitats, with only 3 sampling locations in existing forest stands and 26 in restoration plots. Whilst the random intercepts in the statistical models effectively accommodated this repeated-measure design, incorporating more forest sample points would provide greater statistical power for habitat comparisons. Secondly, the two recording devices (AudioMoths and Song meters) had a significant covariate effect across all fitted models, so detection disparities due to hardware must be considered when interpreting long-term datasets. Finally, positive *Saccopteryx bilineata* detections accounted for only 1.33% of the entire dataset (2,098 positive files across 157,335 recordings), and, as a single-species classifier, cannot measure biodiversity metrics such as species richness or evenness; therefore, results cannot be generalised to neotropical bat assemblages.

### Implications and Direction For Future Studies

Extracting meaningful ecological data from hundreds of hours of recordings can provide important insights into animal behaviour, communication, and spatial occurrence; establishing metrics for assessing habitat recovery and directing restoration efforts (Sharma et al., 2023). The benefits of collecting low-cost, non-invasive acoustic data are constrained by the time-consuming process of manual validation and expert annotation (Brinkløv et al., 2023). Advancements in artificial intelligence offer an effective and efficient solution by automating labour-intensive tasks, thereby reducing workload (Runkel et al., 2021). This study demonstrated the practical advantages of using an automated convolutional neural network to detect a target species in a large acoustic dataset (n=157,336, ∼1,008.7 hours) and the benefits this offers for data processing bottlenecks.

Future bioacoustics monitoring frameworks could incorporate lightweight neural network architectures and expand them to capture multiple tropical bat species for species assemblage surveys. Neotropical bats are an excellent candidate for future studies, as they occupy several dietary niches and provide numerous ecosystem services. Bats are especially effective targets for acoustic monitoring as they continuously vocalise when echolocating. One ecosystem service that insectivorous bats provide is pest control, which is especially relevant in restoration, as insect herbivory damages leaves, affecting the growth and survival of tropical plants (Kelling et al., 2026; Morrison & Lindell, 2012). Evaluating how bat activity correlates with reduced herbivory offers an opportunity for future restoration projects to quantify the functional and economic benefits.

Beyond acoustics, seasonality and meteorological data, such as rainfall, should be considered in future research (Gallacher et al., 2021), as weather and seasons can alter foraging behaviour. With increasing global temperatures, the effects on bat reproductive cycles could be investigated using bioacoustics to locate maternity colonies of cryptic species (Linton & Macdonald, 2018; O’Malley et al., 2023). Furthermore, pairing LiDAR drone technology with acoustic methods could capture high-resolution geospatial data and investigate whether canopy cover and height influence bat activity (Froidevaux et al., 2016).

## Conclusion

Greater sac-winged bats (*Saccopteryx bilineata)* are widely distributed across Brazil, and although they are classified as Least Concern on the IUCN Red List (IUCN, 2015), they provide an effective indicator taxon for evaluating habitat recovery. Understanding how tropical bats respond to restored landscapes in the Arc of Deforestation in the Amazon is paramount to addressing the current biodiversity crisis and understanding how fauna returns to human-degraded landscapes. This research offers encouraging evidence for reforestation, as increased time since restoration positively influenced *Saccopteryx bilineata* occurrence, and habitat also had a strong effect size. These findings highlight the computational efficacy of lightweight 2D-CNNs for efficient data processing of large-scale acoustic studies and tropical reforestation actively restores functional ecological niches for aerial insectivorous bats.

## Acknowledgments

I would like to acknowledge and thank my supervisor, Cristina Leite-Banks. I would also like to extend my thanks to the members of the CaLE lab, especially Rhys and Xiuhan, for collecting the data. Finally, I would also like to acknowledge the use of computational resources, specifically the High Performance Computing (HPC) clusters, provided by the Imperial College Research Computing Service (Harvey, 2017)(http://doi.org/10.14469/hpc/2232).

## Data and Code Availability

### Data availability

Field data recordings used in this thesis are stored on Imperial College London’s Remote Data Storage. Access can be provided upon request from Professor Cristina Bank-Leite, Department of Life Sciences, Silwood Park.

### Code availability-

The 2D-CNN model and analysis scripts are publicly available on GitHub here: https://github.com/secord-stars/Saccopteryx-bilineata-response-to-restoration

### Audio library

The call library used to train the 2D-CNN model is available on Dropbox here: https://www.dropbox.com/scl/fo/awv8ljmxz7hks8ya202cu/AJ3vtsHMxa7veXOviIw v720?rlkey=xr0655r3lvwj9hd26dmhonjsk&st=trbpbuz1&dl=0

IUCN Red List and IGBE map data used in figures are available through their references.

## Generative AI Use Statement

Grammarly Pro was used for spelling and grammatical error checking, and Gemini Pro 3.1 extended thinking was used for troubleshooting and debugging Python code. All written content is original and the authors’ own, and AI-generated content has not been used.

## Appendix

### Supplementary Figures And Tables

Formulas for machine learning performance metrics are calculated using the following: True Positives (TP) and True Negatives. (TN), False Positives (FP) and False Negatives (FN)

**STable 1.** A summary table of the five model performance metrics monitored in model training.

|  |  |
| --- | --- |
| *accuracy* | $\frac{TP + TN}{TP + TN + FP + FN}$ |
| *precision* | $\frac{TP}{TP + FP}$ |
| *recall* | $\frac{TP}{TP + FN}$ |
| *specificity* | $\frac{TN}{TN + FP}$ |
| *F1 score* | $2 \times \frac{\text{precision} \times \text{recall}}{\text{precision} + \text{recall}}$ |

**SFigure 9.**
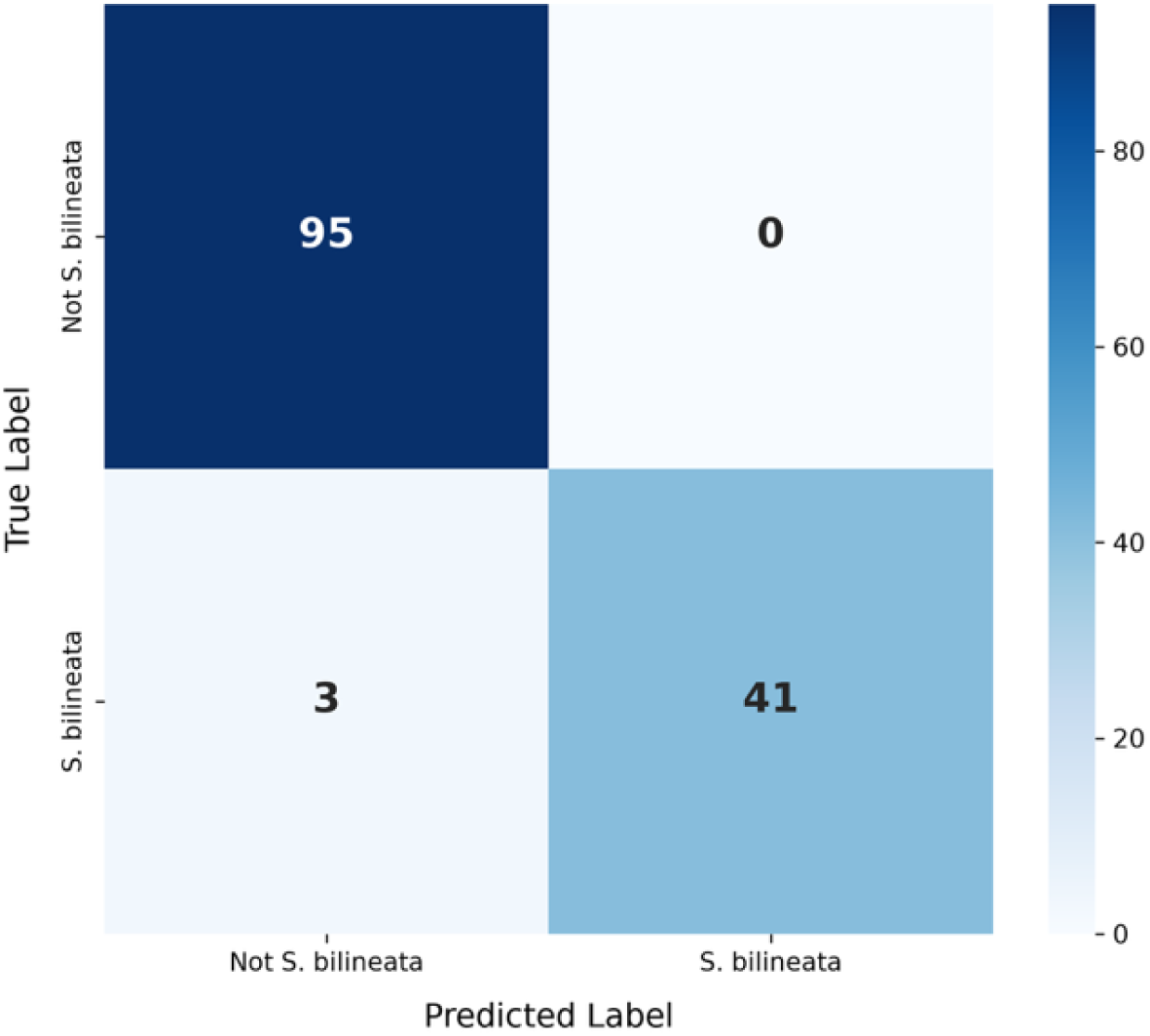
Confusion matrix of the model’s performance on the held-out annotated test set in training. 0 False positives detected, 3 false negatives, n = 139, positives = 44, negatives = 95

**SFigure 10.**
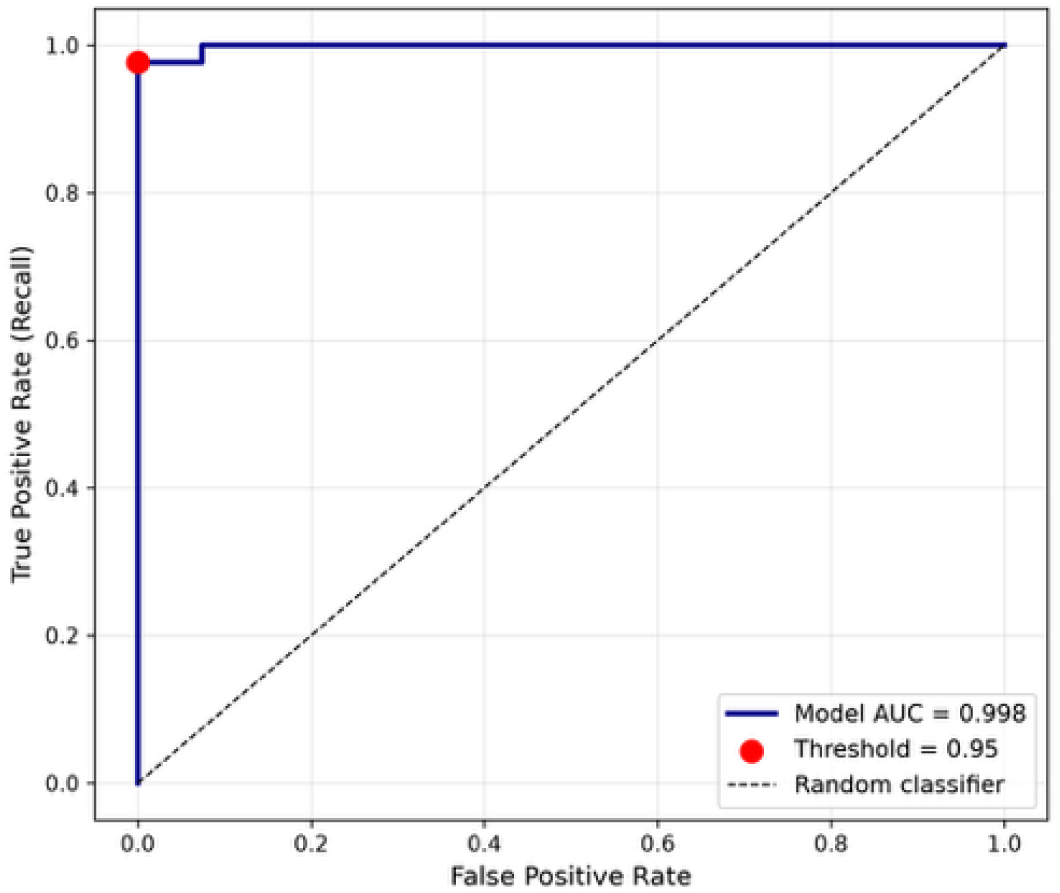
Receiver Operating Characteristic (ROC) curve that shows the binary classification models performance at the classification threshold, t=0.95.

**SFigure 11.**
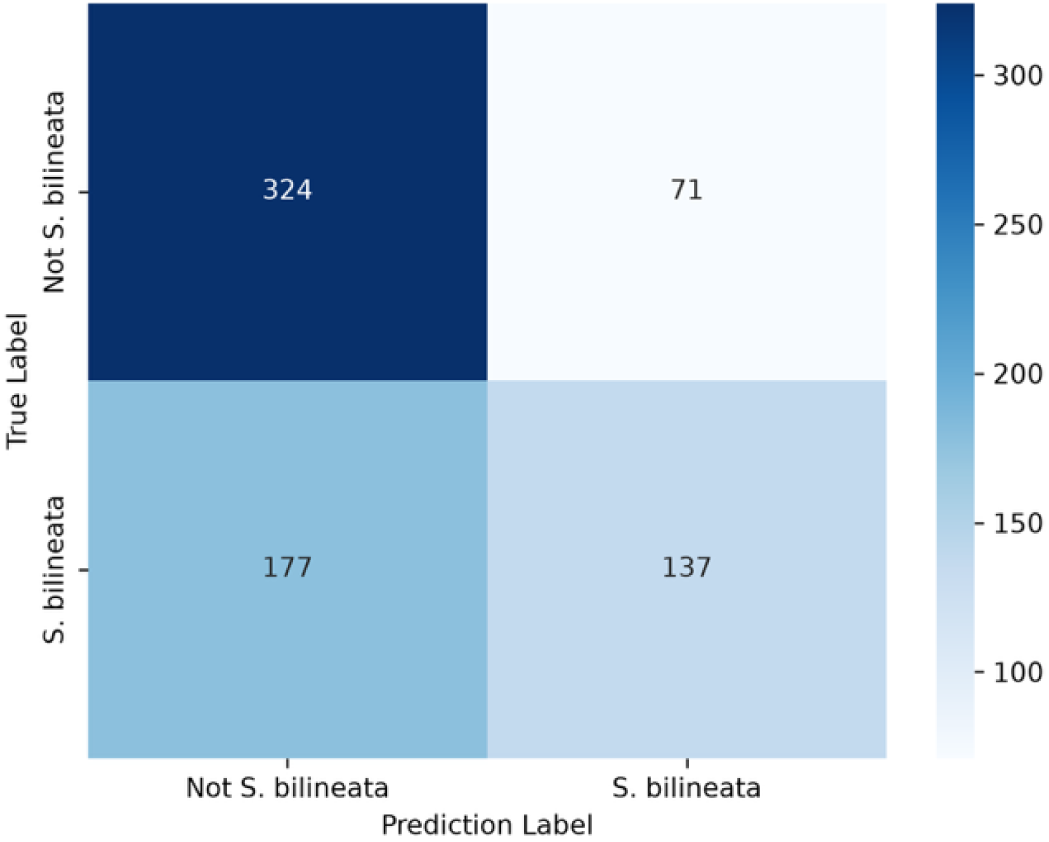
Confusion matrix of the stratified validation audi on the non-annotated dataset, n = 709 files, positives = 208, negatives = 501 . Overall accuracy 65.02%, precision 65.87%, recall 43.63%.

**SFigure 12.**
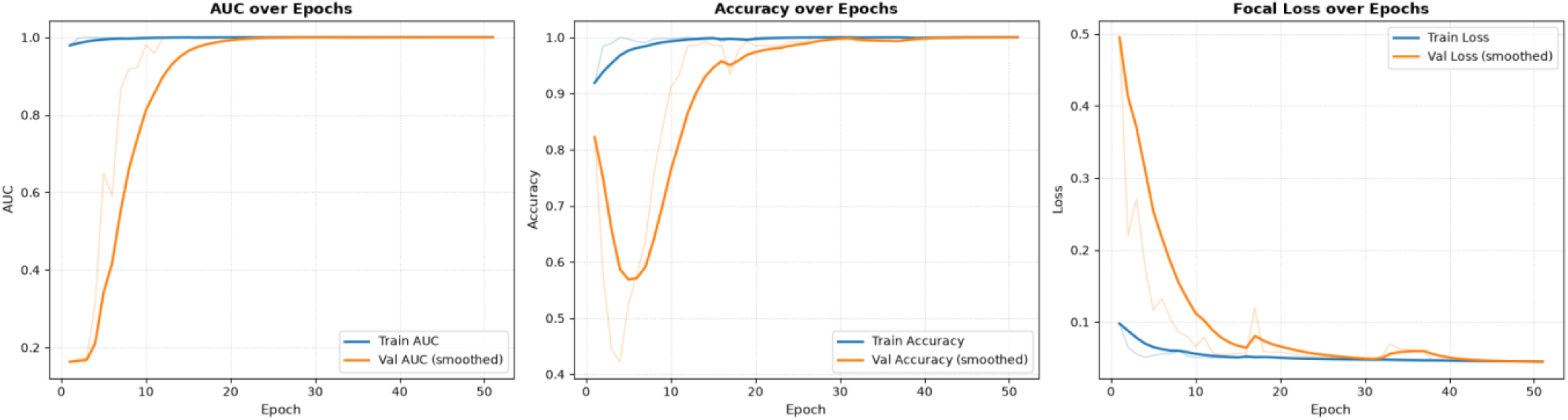
Performance metrics across training epochs. a) Area under the receiver operating characteristic curve (ROC-AUC) b) Overal classification accuracy c) Overall focal loss. Blue lines show training set performance and orange show validation set performance

**SFigure 13.**
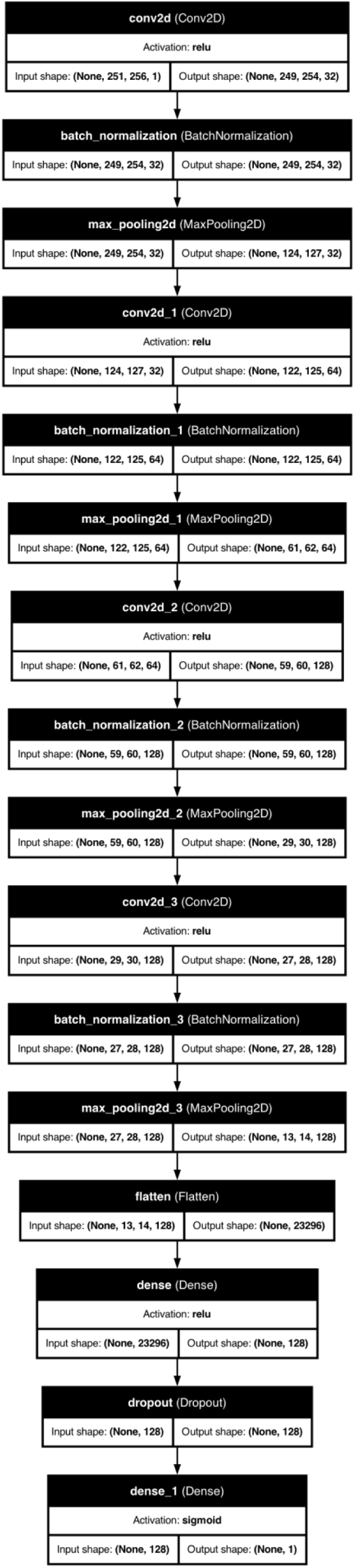
A 2D flowchart of the model architecture, generated using visual.keras in Python. The cells show the input and output tensor shapes and how the input is flattened and collapsed in the final dense layer with a sigmoid activation

**SFigure 14.**
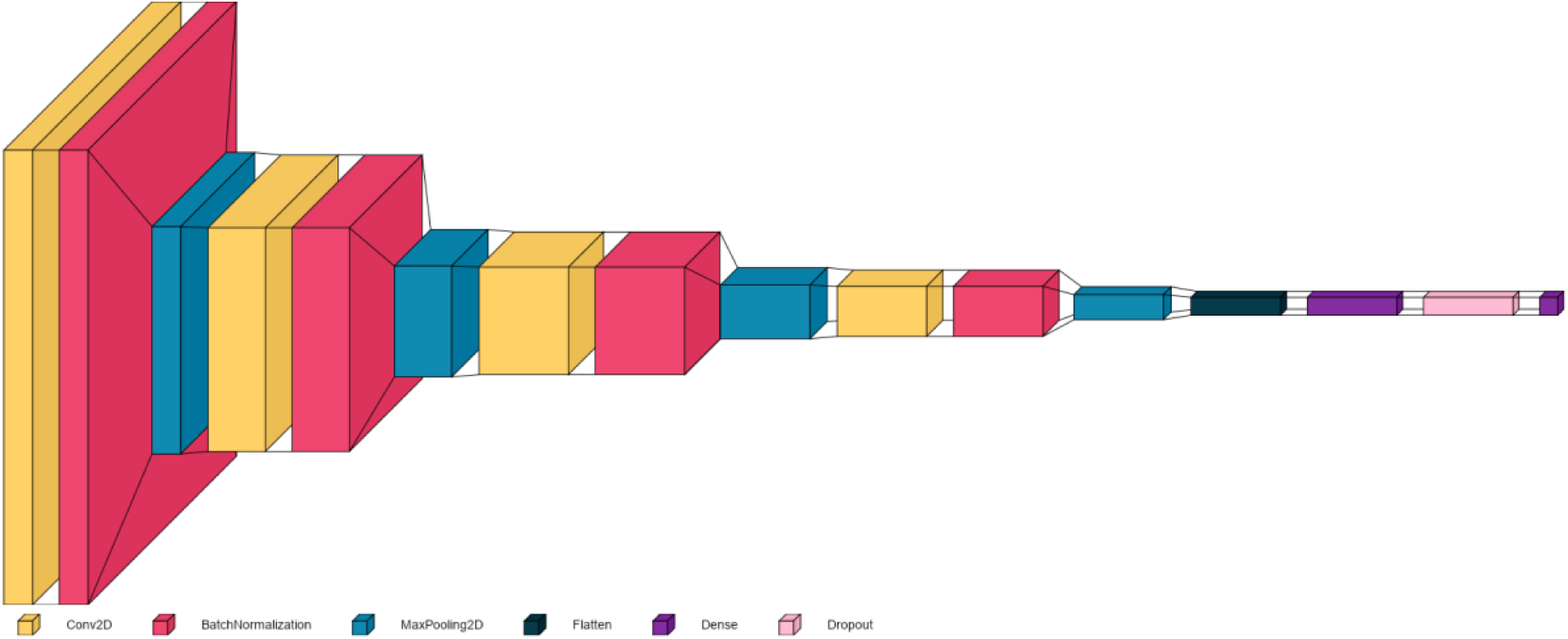
A 3D block map of the model architecture, generated using visual. keras in Python. This visualisation shows the four layers of the model, with a single layer represented by a yellow, red and blue cubes. The size of the cubes reflects the tensor shapes and shows how the data is pooled, before being flattened and collapsed to a single output in the final dense layer.

**STable 2.** A summary table of the sampling point, habitat, replanting date, deployment and collection date across the three sampling years 2023, 2024, 2025

| Sample Point | Habitat | Date planted | Deployment Date 2023 | Collection Date 2023 | Deployment Date 2024 | Collection Date 2024 | Deployment Date 2025 | Collection Date 2025 |
| --- | --- | --- | --- | --- | --- | --- | --- | --- |
| TU001 | Restoration | 17/02/2025 | 10/02/2023 | 16/02/2023 | 07/03/2024 | 16/03/2024 | 12/03/2025 | 19/03/2025 |
| TU002 | Restoration | 17/02/2025 | 08/02/2023 | 14/02/2023 | 06/03/2024 | 29/03/2024 | 12/03/2025 | 26/03/2025 |
| TU003 | Restoration | 12/02/2025 | — | — | 08/03/2024 | 18/03/2024 | 13/03/2025 | 16/03/2025 |
| TU004 | Restoration | — | 07/02/2023 | 14/02/2023 | 22/03/2024 | 29/03/2024 | 23/03/2025 | 26/03/2025 |
| TU005 | Restoration | 19/02/2025 | 09/02/2023 | 15/02/2023 | 14/04/2024 | 21/04/2024 | 22/03/2025 | 29/03/2025 |
| TU006 | Restoration | 23/05/2023 | 10/02/2023 | 10/02/2023 | 20/03/2024 | 30/03/2024 | 12/03/2025 | 19/03/2025 |
| TU007 | Restoration | 31/08/2023 | 08/02/2023 | 14/02/2023 | 21/03/2024 | 29/04/2024 | 13/03/2025 | 27/03/2025 |
| TU008 | Restoration | 21/01/2025 | 08/02/2023 | 14/02/2023 | 15/04/2024 | 21/04/2024 | 22/03/2025 | 29/03/2025 |
| TU009 | Restoration | 11/11/2024 | — | — | 20/03/2024 | 28/03/2024 | 11/03/2025 | 18/03/2025 |
| TU010 | Restoration | 14/01/2025 | 08/02/2023 | 14/02/2023 | 14/04/2024 | 21/04/2024 | 22/03/2025 | 28/03/2025 |
| TU011 | Restoration | 09/08/2023 | 17/02/2023 | 20/02/2023 | 21/03/2024 | 29/03/2024 | 13/03/2025 | 26/03/2025 |
| TU012 | Restoration | 11/09/2024 | 09/02/2023 | 16/02/2023 | 15/04/2024 | 22/04/2024 | 22/03/2025 | 28/03/2025 |
| TU013 | Restoration | — | 08/02/2023 | 15/02/2023 | 13/04/2024 | 21/04/2024 | 23/03/2025 | 26/03/2025 |
| TU014 | Restoration | 23/11/2023 | 08/02/2023 | 15/02/2023 | 14/04/2024 | 22/04/2024 | 21/03/2025 | 29/03/2025 |
| TU015 | Restoration | 23/12/2023 | 08/02/2023 | 15/02/2023 | 22/03/2024 | 30/03/2024 | 13/03/2025 | 17/03/2025 |
| TU016 | Restoration | 30/11/2023 | 08/02/2023 | 15/02/2023 | 13/04/2024 | 21/04/2024 | 13/03/2025 | 17/03/2025 |
| TU017 | Restoration | 30/10/2023 | 08/02/2023 | 15/02/2023 | — | — | 13/03/2025 | 19/03/2025 |
| TU018 | Restoration | 05/04/2024 | 09/02/2023 | 16/02/2023 | 15/04/2024 | 22/04/2024 | 21/03/2025 | 31/03/2025 |
| TU019 | Restoration | 13/12/2023 | — | — | 14/04/2024 | 21/04/2024 | 23/03/2025 | 31/03/2025 |
| TU020 | Restoration | 29/04/2023 | 15/02/2023 | 23/02/2023 | 20/03/2024 | 29/03/2024 | 11/03/2025 | 19/03/2025 |
| TU021 | Restoration | 13/12/2023 | 07/02/2023 | 15/02/2023 | 14/04/2024 | 21/04/2024 | 22/03/2025 | 28/03/2025 |
| TU022 | Restoration | 31/05/2023 | 15/02/2023 | 23/02/2023 | 20/03/2024 | 29/03/2024 | 12/03/2025 | 26/03/2025 |
| **TU023** | Restoration | 07/11/2023 | 08/02/2023 | 15/02/2023 | 13/04/2024 | 21/04/2024 | 20/03/2025 | 30/03/2025 |
| **TU024** | Restoration | 0/08/2024 | 08/02/2023 | 15/02/2023 | 14/04/2024 | 22/04/2024 | 21/03/2025 | 28/03/2025 |
| **TU025** | Restoration | 19/05/2023 | — | — | 20/03/2024 | 29/03/2024 | 12/03/2025 | 20/03/2025 |
| **TU026** | Restoration | 12/07/2023 | 08/02/2023 | 15/02/2023 | 21/03/2024 | 30/03/2024 | 24/03/2025 | 28/03/2025 |
| **TU027** | Forest | — | 08/02/2023 | 15/02/2023 | 22/03/2024 | 30/03/2024 | 23/03/2025 | 27/03/2025 |
| **TU028** | Forest | — | 10/02/2023 | 17/02/2023 | 22/03/2024 | 30/03/2024 | 12/03/2025 | 18/03/2025 |
| **TU029** | Forest | — | 10/02/2023 | 17/02/2023 | 14/04/2024 | 16/04/2024 | 12/03/2025 | 17/03/2025 |

